# Vigour/tolerance trade-off in cultivated sunflower (*Helianthus annuus*) response to salinity stress is linked to leaf elemental composition

**DOI:** 10.1101/447128

**Authors:** Andries A. Temme, Kelly L. Kerr, Lisa A. Donovan

## Abstract

Developing more stress-tolerant crops will require greater knowledge of the physiological basis of stress tolerance. Here we explore how biomass declines in response to salinity relate to leaf traits across twenty genotypes of cultivated sunflower (Helianthus annuus). Plant growth, leaf physiological traits, and leaf elemental composition were assessed after 21 days of salinity treatments (0, 50, 100, 150, or 200 mM NaCl) in a greenhouse study. There was a trade-off in performance such that vigorous genotypes, those with higher biomass at zero mM NaCl, had both a larger absolute decrease and proportional decrease in biomass due to increased salinity. More vigorous genotypes at control were less tolerant to salinity. Contrary to expectation, genotypes with a low increase in leaf Na and decrease in K:Na were not better at maintaining biomass with increasing salinity. Rather, genotypes with a greater reduction in leaf S and K content were better at maintaining biomass at increased salinity. While we found an overall trade-off between sunflower vigour and salt tolerance, some genotypes were more tolerant than expected. Further analysis of the traits and mechanisms underlying this trade-off may allow us to breed these into high vigour genotypes in order to increase their salt tolerance.

## Introduction

Human population levels are predicted to reach 9.7 billion by the year 2050 (UN DESA 2015), which will apply pressure on food-production systems in order to keep pace with increased demand due to population growth (McCouch et al., 2013). Moreover, global shifts in diet toward foodstuffs that are more land-intensive to produce will place additional pressure on agricultural production (Kastner, Rivas, Koch, & Nonhebel, 2012). To increase crop productivity and improve food security for the 21st century, it will be necessary for food production to occur on less ideal cropland and under more stressful growing conditions (Godfray, 2011; Tilman, Balzer, Hill, & Befort, 2011). Salinity is estimated to affect ca. 20% to 50% of irrigated land due to salt accumulation in the soil from poor irrigation practices or from seawater, resulting in crop yield reductions and even plant death and crop loss (Flowers & Yeo, 1995; Munns, 2002; Munns & Tester, 2008). To allow for higher productivity on salinized lands there is a need for the development of more salt tolerant crops. The development of salt-tolerant crops will require greater knowledge on the physiological basis of salt tolerance.

In the face of abiotic stresses such as high salinity, modern crops are generally thought to exhibit reduced stress tolerance as compared to their wild progenitors (Mayrose et al. 2011, Koziol et al. 2012). While there are many ways to define tolerance (Deinlein et al., 2014; Tester & Langridge, 2010; Vinocur & Altman, 2005), here we refer to greater tolerance as a lower proportional effect of stress. A reduced tolerance in modern crops suggests that the capacity to tolerate stress has been lost during domestication (Tanksley & McCouch, 1997). Furthermore, there could be trade-offs between high productivity under unstressed conditions (high vigour, a key feature of crops compared to their wild relatives) and tolerance to stress (Mayrose, Kane, Mayrose, Dlugosch, & Rieseberg, 2011). Thus a key goal for the future is to determine the extent of and reduce these trade-offs (Sadras & Richards, 2014) in order to have both a highly productive and stress-tolerant crop variety.

Salinity, a stress impacting productivity, manifests in plants as both an osmotic stress due to the effect of salt on soil water potential, and as an ionic stress due to the accumulation of potentially toxic sodium (Na^+^) ions (Arzani & Ashraf, 2016; Munns, 2011). An increase in soil salinity inhibits the ability of plants to uptake soil water, and increased concentrations of salt ions in the plant tissue can impair metabolic processes and photosynthesis (Mäser, Gierth, & Schroeder, 2002). Genotypes tolerant to these osmotic and ionic stresses may have different mechanisms to cope with them, such as reduced stomatal conductance to conserve water (Munns & Tester, 2008) and differential exclusion or sequestration of Na^+^ through the plant (Deinlein et al., 2014; Schroeder et al., 2013). Thus a key step towards improving salt tolerance is identifying trade-offs between plant growth and salt tolerance across genotypes and then measurement of associated traits that suggest which of these tolerance mechanisms are employed. This information can then form the basis of more mechanistic studies of greater tolerance such as strong root membrane potentials to keep Na^+^ out (Bose et al., 2015; Chen et al., 2007), and sequestration of or through sequestering Na^+^ in the vacuole (Ballesteros, Blumwald, Pedro Donaire, & Belver, 1997; Blumwald, 2000).

Compared to other crops species, cultivated sunflower has been shown to be moderately salt tolerant (Katerji, van Hoorn, Hamdy, & Mastrorilli, 2000), as has safflower, a closely related crop species (Yeilaghi, Arzani, & Ghaderian, 2015). Additionally, cultivated sunflower shows genotypic variation in response to abiotic stresses, including drought (eg. Ahmad et al. 2009), nutrients (eg. Cechin and de Fatimas Fumis 2004), and salinity (eg. (G. Ceccoli et al., 2015; Katerji et al., 2000; Rawson & Munns, 1984; Shi & Sheng, 2005). Given the moderate salt tolerance and putative genotypic variation in response to salinity in sunflower, there is a potential for identifying a range of salinity tolerances in cultivated sunflower genotypes that could be linked to physiological mechanisms underlying salt tolerance given a broad set of genotypes. However, contrary to several other important crops (Flowers, 2004), there are relatively few studies that have studied sunflower salinity tolerance across a range of salinity levels and a large number of genotypes, which has made it difficult to identify trade-offs across the sunflower germ plasm pool and general mechanisms of tolerance.

For cultivated sunflowers, studies of relatively small number of genotypes and limited soil salinity treatments (have provided some insights into tolerance mechanisms (e.g. (Akram, Ashraf, & Akram, 2009; Ashraf, 1999; Delgado & Sanchez-Raya, 1999; Shi & Sheng, 2005; Sohan, Jasoni, & Zajicek, 1999; Torabian, Zahedi, & Khoshgoftar, 2016, 2017). To mitigate the osmotic stress imposed by soil salinity, sunflowers use mechanisms to reduce water loss while maximizing water uptake, including a reduction in leaf area (Rawson & Munns, 1984; Steduto, Albrizio, Giorio, & Sorrentino, 2000) and osmotic adjustment (Deinlein et al., 2014). Other mechanisms specifically mitigate ion toxicity effects, such as limiting Na^+^ uptake (Mutlu & Bozcuk, 2005), discrimination between potassium (K^+^) and Na^+^ (Shabala & Cuin, 2008), and reducing cytoplasmic Na^+^ concentrations through dilution, excretion or sequestration (Munns, 2002). The exploration of a more comprehensive network of correlated trait shifts (Poorter, Anten, & Marcelis, 2013), including nutrients beyond Na^+^ and K^+^, across a range of salinity concentrations and genotypes may shed more light on sunflower’s response to salinity and provide directions for further research for this important oil seed crop.

Here, we examined the response of growth and functional traits, including elemental composition, of twenty cultivated lines of sunflower (*Helianthus annuus L*.) under a wide range of salinity concentrations. We asked the following questions: **1)** What is the decline in growth over a range of salinity concentrations, and is there evidence for differential responses among genotypes?; **2)** Is there evidence for a trade-off between growth under unstressed (vigour) versus stressed conditions? **3)** How do plant traits adjust with changes in soil salinity, and are they correlated with the effect of salinity on growth performance?

## Materials and Methods

### Study Design

Twenty inbred genotypes (**Supplemental table 1**), including elite varieties and landraces, of cultivated sunflower (*Helianthus annuus L*.) were selected from a diversity panel, sunflower association mapping (SAM) population of 288 genotypes (Mandel, Dechaine, Marek, & Burke, 2011; Nambeesan et al., 2015). Genotypes were selected based on their differential responses in previous abiotic stress studies (Bowsher et al., 2017; Masalia, Temme, Torralba, & Burke, 2018). Plants were grown in a split plot design with four replicates per treatment and genotype at the Plant Biology Greenhouse on the University of Georgia campus located in Athens, GA from September to October of 2016.

Achenes (hereafter “seeds”) for all 20 genotypes were planted on September 12^th^ 2016 into seedling trays with a soil medium composed of a 3:1 ratio of sand to Turface MVP^®^ (Turface Athletics, PROFILE Products, LLC, Buffalo Grove, IL). Each seed was placed into a 2 mm depression, covered with soil, and treated with a 0.45 g/L solution of a broad-spectrum fungicide to inhibit fungal growth (Banrot, Everris NA Inc., Dublin, OH). Seedlings were transplanted into 5L plastic pots four days after planting and watered daily until treatment initiation.

Twenty plastic-lined ponds were constructed, with five ponds placed on each of four greenhouse benches. The five ponds on each bench were randomly assigned a salinity treatment of either 0, 50, 100, 150 and 200 mM sodium chloride (NaCl). Twenty, 30 cm tall, 5L pots with transplanted seedlings were placed into each pond with the bottom 8-10 cm of the pot standing in water, totaling 100 pots per bench and 400 pots across all benches. Each pot was filled with the same soil medium used in the seedling trays. Pots also received 40 g 15-9-12 (N-P-K) Osmocote Plus blend (Osmocote, The Scotts Company, Marysville, OH), and supplemental calcium in the form of 5 ml of gypsum (Performance Minerals Corporation, Birmingham, AL) and 5 ml of garden lime powder (Austinville Limestone, Austinville, VA). The top 10 cm of soil was well-mixed to ensure an even distribution of these amendments.

Treatments were initiated nine days after planting. The appropriate treatment solution (0, 50, 100, 150 or 200 mM NaCl) was added to each pond which inundated the lower 8-10 cm of the pots. The first day, the solution in each pond was allowed to infiltrate the soil from the bottom of the pot in order to reduce salinity shock in the seedlings. Pots were top-watered daily for the following three days with 500 mL of solution from the corresponding ponds to homogenize the salinity in the soil and the pond. During this interval, salinity concentrations of the treatment solutions were checked daily with an electric conductivity (EC) probe (HI 8733, Hanna Instruments Inc., Woonsocket, USA) and a salinity refractometer (Reichert Technologies, Munich, Germany), with fresh or salt water added as needed to reestablish the desired concentration. Top-watering was then discontinued unless the soil appeared dry and there was no visible moisture 2 cm below the soil surface. Plants were harvested after 21 days of treatment.

### Measurements

Height from the base of the stem to the tip of the apical meristem was measured to the nearest 0.5 cm on all plants at 7,14, and 21 days after treatment initiation. Stem diameter was measured at the base of the stem only at 21 days after treatment initiation to avoid damaging the developing plants. Relative height growth (Rel. ht. gr.) was calculated for for all plants using the equation: Rel. ht. gr. = (*In (H_1_-ln (H_2_))/t_2_-t_1_*, where *ln* is natural logarithm, *H_2_* is plant height at time two, *H_1_* is plant height at time one, *t_2_* is time two, and *t_1_* is time one. The relative height growth was determined by averaging relative height growth for each interval between the three time points at which plant height was measured. The mean relative height growth (rel. ht. gr.) value was used for making comparisons within and across genotypes.

Quantum yield (QY) was measured using a chlorophyll fluorometer (FluorPen, Photon Systems Instruments, Drásov, Czech Republic) 17 days after treatment onset, at predawn (0400 h - 0600 h) and midday (1200 h - 1400 h). Two readings were taken on the most recent fully expanded leaf (MRFEL) from each plant, one on each side of the leaf midrib, and the average of the two readings was rounded to the nearest 0.1 unit. Chlorophyll concentration was non-destructively measured at harvest using a chlorophyll concentration meter (MC-100, Apogee Instruments, Inc., Logan, UT). Two readings were taken on the MRFEL, one on each side of the leaf midrib, and the average of the two readings was rounded to the nearest 0.1 CCI (chlorophyll concentration index).

Living plants were harvested for biomass 21 days after treatment onset. Biomass was separated into the most-recently-fully-expanded-leaf (MRFEL), remaining leaf tissue, stem, and roots. Roots were washed on a 2 mm wire screen and gently squeezed to remove excess water. All biomass tissue samples were dried at 60 °C for 48 hrs. After drying, leaf, stem and reproductive bud samples were weighed to the nearest 0.01 g. Root samples were weighed to the nearest 0.0001 g. Total biomass was determined by summing all tissue types. Root mass fraction (RMF), leaf mass fraction (LMF) and shoot mass fraction (SMF) were calculated by dividing the root, leaf and shoot biomass values by the total biomass values for each plant sample, respectively.

At harvest, the removed MRFEL (leaf and petiole) was placed onto a flatbed scanner (Canon CanoScan LiDE120) and scanned as a 300 dpi JPG image. The MRFEL was then dried at 60 °C for 48 hrs. After drying, the MRFEL was separated from its petiole, and both MRFEL leaf and MRFEL petiole were weighed to the nearest .0001 g. Leaf scans were processed using ImageJ (NIH, USA, http://rsb.info.nih.gov/ii/) by converting scans to binary and counting the number of pixels in leaf blade and petiole. Specific leaf area (SLA mm^2^/g) was then calculated by dividing the leaf blade area by the leaf blade weight.

### Ion Analysis

The dried MRFEL samples were bulked by genotype and treatment, resulting in four MRFEL samples per genotype per treatment. Bulk MRFEL samples, without petioles, were ground into powder using a Wiley Mill and a Qiagen tissuelyser (Qiagen, Venlo, Netherlands) with a steel bead. This yielded too little tissue to analyse both foliar nitrogen and other element concentrations, so the bulk MRFEL powder was used to determine only leaf nitrogen content. We ground all other leaf tissue, excluding petioles, similar to the MRFEL samples to determine the other element concentrations. MRFEL ion concentration and rest of leaves ion concentration were highly correlated (**Supplemental figure 1**).

Powder from each genotype and treatment combination was placed into a 2 ml Eppendorf tube and shipped for nitrogen analysis and Inductively Coupled Argon Plasma Optical Emission (ICP) Analysis (Midwest Laboratories, Omaha, NB). Analyses provided total element concentrations of the leaf tissues for the following elements: nitrogen (N) via the Dumas method, and phosphorus (P), potassium (K), magnesium (Mg), calcium (Ca), sulfur (S), sodium (Na), iron (Fe), manganese (Mn), boron (B), copper (Cu), zinc (Zn) via ICP analysis.

#### Statistical Analysis

Linear mixed effects models were used to fit the relationships between treatment (0, 50, 100, 150, 200 mM NaCl) and values of the growth and physiological response variables (see Table 1) across all genotypes. To account for the split plot design, pond was nested within bench and was treated as a random factor. Linear models were used to fit the relationships between treatment and mean values of the nitrogen and ICP elemental concentrations (N, P, K, Mg, Ca, S, Na, Fe, Mn, B, Cu, Zn) across all genotypes. Because these samples were bulked, the effect of bench could not be determined. The assumptions of these models were checked by examining plots of the residuals. Genotype mean trait values at each salinity treatment were calculated as estimated marginal means using the R package ‘emmeans’ (Lenth, 2018). Statistical analyses were conducted in R version 3.2.3 (R Core Team 2015) using mixed models (‘lme4’ package; Bates 2014). Estimates for the effect of genotype, salinity treatment, and their interaction were made by Walds Analysis of Deviance Type 3 Anova using a chi-squared (□^2^) test (‘car’ package; (Fox & Weisberg, 2011).

**Table 1.** Median trait value and range of twenty sunflower genotypes in five salinity treatments (0-200 mM NaCl). Table shows median trait values (and range of trait values between genotypes in brackets) for morphological, physiological and chemical composition traits across estimated marginal means of all twenty sunflower genotypes. Vigour-related traits (mass, size) were additionally natural log transformed to account for allometry in estimating the proportional effect of salinity on trait values. Asterisks denote p-value of NaCl treatment (T), genotype differences (G) or their interaction (G*T). [.=p<0.1, *=p<0.05, **=p<0.01, ***=p<0.001]. Abbreviations: SLA, specific leaf area (m^2^_leaf_ g_leaf_^−1^); LAR, leaf area ratio (m^2^_leaf_ g_plant_^−1^); SSL, specific stem length (cm_stem_ g_stem_^−1^); Rel. ht. gr, relative height growth (cm_stem_ cm_stem_^−1^ day^−1^); ns, no significance.

| Trait | 0 mM | 50 mM | 100 mM | 150 mM | 200 mM | Treatment (T) | Genotype (G) | G*T |
| --- | --- | --- | --- | --- | --- | --- | --- | --- |
| Total biomass (g) | 1.5 (0.43-4.18) | 0.97 (0.42-2.44) | 0.72 (0.32-2.07) | 0.44 (0.2-1.05) | 0.52 (0.36-0.9) | *** | *** | *** |
| Ln(Total biomass (g) ) | 0.34 (-0.58-1.39) | -0.13 (-1.17-0.72) | -0.45 (-1.08-0.69) | -0.87 (-1.57--0.07) | -0.83 (-1.24--0.26) | *** | *** | ns |
| Leaf mass (g) | 0.87 (0.33-2.28) | 0.5 (0.22-1.51) | 0.37 (0.16-1.1) | 0.28 (0.14-0.63) | 0.33 (0.21-0.51) | *** | *** | *** |
| Ln(Leaf mass (g) ) | -0.17 (-0.98-0.79) | -0.79 (-1.92-0.25) | -1.08 (-1.78-0.05) | -1.33 (-2.03--0.57) | -1.28 (-1.7--0.75) | *** | *** | ns |
| Root mass (g) | 0.23 (0.11-0.42) | 0.21 (0.07-0.36) | 0.15 (0.07-0.32) | 0.08 (0.03-0.24) | 0.1 (0.02-0.2) | ** | *** | * |
| Ln(Root mass (g) ) | -1.56 (-2.34--0.89) | -1.67 (-3.31--1.11) | -2.08 (-2.89--1.12) | -2.6 (-3.47--1.38) | -2.55 (-4.06--1.71) | ** | *** | ns |
| Stem mass (g) | 0.35 (0-1.52) | 0.19 (0.04-0.65) | 0.15 (0.04-0.65) | 0.09 (0.03-0.3) | 0.08 (0.03-0.27) | *** | *** | *** |
| Ln(Stem mass (g) ) | -1.2 (-2.73-0.35) | -1.77 (-2.85--0.79) | -1.93 (-3.12--0.5) | -2.41 (-3.39--1.24) | -2.54 (-3.74--1.45) | *** | *** | ns |
| Plant height (cm) | 22.06 (2.82-56.75) | 16.13 (4.21-45.29) | 14.25 (4.5-41.33) | 10.43 (3.08-26.56) | 9.96 (1.97-26.97) | *** | *** | *** |
| Ln(Plant height (cm) ) | 3.06 (1.62-4.03) | 2.76 (1.66-3.74) | 2.62 (1.5-3.68) | 2.33 (1.23-3.26) | 2.26 (0.89-3.3) | *** | *** | ns |
| Rel. ht gr ($cm\ cm^{-1}\ d^{-1}$) | 0.1 (-0.03-0.12) | 0.07 (0.05-0.1) | 0.07 (0.04-0.08) | 0.05 (0.01-0.07) | 0.04 (-0.03-0.08) | *** | *** | ** |
| Diameter (mm) | 5.29 (3.34-8.56) | 3.93 (2.88-6.57) | 3.49 (2.66-5.91) | 3.16 (2.37-4.68) | 3.25 (2.05-4.28) | *** | *** | *** |
| SLA ($m^2\ g\ leaf^{-1}$) | 45.31 (32.55-48.31) | 36.22 (27.31-43.65) | 33.27 (26.78-39.9) | 29.83 (24.6-32.91) | 28.9 (19.95-38.93) | *** | *** | ns |
| LAR ($m^2\ g\ plant^{-1}$) | 28.3 (17.9-32.31) | 20.46 (13.08-24.89) | 19.33 (14.04-23.19) | 18.56 (13.75-21.15) | 18.01 (13.39-23.5) | *** | *** | * |
| Leaf mass fraction | 0.62 (0.51-0.69) | 0.56 (0.43-0.65) | 0.59 (0.43-0.66) | 0.62 (0.51-0.68) | 0.62 (0.54-0.67) | ns | *** | ns |
| Root mass fraction | 0.16 (0.09-0.26) | 0.21 (0.13-0.31) | 0.19 (0.13-0.31) | 0.17 (0.13-0.29) | 0.2 (0.13-0.3) | * | *** | ns |
| Stem mass fraction | 0.23 (0.12-0.36) | 0.22 (0.09-0.38) | 0.22 (0.1-0.44) | 0.2 (0.15-0.31) | 0.19 (0.04-0.33) | . | *** | ns |
| SSL ($cm\ g\ stem^{-1}$) | 76.84 (40.94-129.79) | 103.21 (62.79-152.81) | 105.49 (65-160.22) | 116.01 (53.52-180.56) | 118.16 (85.24-205.53) | ** | ns | ns |
| Chlorophyll index | 14.79 (12.3-24.79) | 17.14 (14.35-31.92) | 19.48 (11.16-30.42) | 16.03 (12.05-28.45) | 17.09 (11.82-30) | ns | *** | *** |
| Quantum yield<sub>dark</sub> | 0.82 (0.8-0.85) | 0.82 (0.8-0.84) | 0.82 (0.8-0.83) | 0.81 (0.76-0.83) | 0.79 (0.59-0.81) | ** | * | *** |
| Quantum yield<sub>light</sub> | 0.7 (0.65-0.76) | 0.69 (0.64-0.75) | 0.67 (0.58-0.69) | 0.66 (0.36-0.74) | 0.61 (0.45-0.75) | * | ns | ** |
| QY.diff | 0.12 (0.08-0.16) | 0.13 (0.08-0.19) | 0.15 (0.12-0.24) | 0.15 (0.05-0.45) | 0.18 (-0.02-0.32) | ns | ns | * |
| QY.avg | 0.76 (0.73-0.8) | 0.75 (0.73-0.79) | 0.74 (0.7-0.76) | 0.74 (0.59-0.76) | 0.71 (0.59-0.78) | ** | ns | *** |
| [Na] (%) | 0.01 (0-0.03) | 0.3 (0.07-0.72) | 1.2 (0.23-2.72) | 2.9 (1.21-4.96) | 4.7 (0.72-7.95) | *** | ns | ** |
| [K] (%) | 5.81 (4.88-6.93) | 5.15 (4.03-6.27) | 4.61 (2.81-6.07) | 4.17 (1.8-5.08) | 3.94 (1.62-5.49) | ns | *** | *** |
| [Na]/[K] (ratio) | 0 (0-0) | 0.06 (0.01-0.17) | 0.26 (0.04-0.82) | 0.68 (0.24-2.11) | 1.14 (0.13-3.84) | * | ns | *** |
| [N]<sub>SLA leaf</sub> (%) | 7.34 (6.59-7.72) | 6.61 (5.93-7.45) | 6.28 (5.07-7.13) | 5.88 (4.62-6.64) | 5.58 (5.12-6.83) | ** | ns | . |
| [S] (%) | 0.59 (0.47-0.78) | 0.5 (0.43-0.58) | 0.47 (0.4-0.55) | 0.47 (0.37-0.68) | 0.5 (0.33-0.6) | ns | * | . |
| [P] (%) | 0.44 (0.34-0.54) | 0.4 (0.29-0.51) | 0.32 (0.25-0.54) | 0.36 (0.2-0.45) | 0.26 (0.14-0.5) | ns | . | * |
| [Mg] (%) | 0.41 (0.29-0.64) | 0.46 (0.33-0.7) | 0.54 (0.35-0.81) | 0.56 (0.34-0.74) | 0.55 (0.37-0.7) | * | *** | * |
| [Ca] (%) | 1.17 (0.92-1.77) | 1.34 (0.84-2.31) | 1.63 (0.99-2.19) | 1.43 (0.97-2.04) | 1.52 (0.71-1.83) | ns | *** | ns |
| [Fe] (ppm) | 126 (91-265) | 165 (88-565) | 150 (71-452) | 101 (66-382) | 102 (57-255) | . | . | ns |
| [Mn] (ppm) | 269 (155-405) | 442.5 (235-1042) | 455.5 (269-1058) | 434 (272-728) | 409.5 (217-656) | ns | ns | ns |
| [Cu] (ppm) | 28 (21-38) | 34 (28-57) | 41.5 (33-56) | 43 (31-66) | 39 (27-68) | ns | ns | ns |
| [Zn] (ppm) | 54 (42-83) | 71 (42-111) | 78.5 (37-125) | 75 (37-104) | 73.5 (38-103) | ns | ** | ns |
| [B] (ppm) | 83.5 (64-119) | 97 (67-135) | 98 (71-138) | 113 (74-150) | 96.5 (60-142) | ** | *** | *** |

Correlation matrices for traits at each salinity level were created by correlating all pairwise trait combinations using pearson correlation. Correlation strength was tested using standardized major axis (SMA) regression (Warton, Duursma, Falster, & Taskinen, 2012) to account for uncertainty in both traits. To explore correlated plasticity in traits, the slope of each genotype’s trait adjustment to increasing salinity was calculated from the mixed model (or linear model for elemental concentrations). These slopes were then correlated and tested as above to determine whether a stronger trait adjustment to salinity in one trait was correlated with a stronger or weaker trait adjustment in another trait. Correlation networks were visualized using the R packages ‘igraph’ (Csardi & Nepusz, 2006) and ‘ggraph’ (Pedersen, 2018). All other graphs were made using the R package ‘ggplot2’ (Wickham, 2009) with twenty distinct colours for all genotypes from ‘https://sashat.me/2017/01/11/list-of-20-simple-distinct-colors/’.

## Results

### Effects of salinity on growth

Out of 35 traits measured, 24 were significantly affected by increasing salinity, 24 differed among genotypes and 19 showed an interaction (G*T) between genotype (G) and the response to salinity (T) **(Table 1)**. All genotypes decreased in height, biomass, and leaf area ratio (LAR). Salinity had a large effect on biomass accumulation, the extent of which differed significantly between genotypes (p<0.001) **(Figure 1a)**. After natural log transformation, we could not detect strong differences between genotypes and the proportional effect of salinity on biomass with increasing salinity **(Figure 1b)**, possibly due to high variance or a limited range in slopes of treatment effect. Using the average slope across all genotypes in Figure 1b, we did determine that for each 50 mM increase in salinity, biomass was reduced by 25% as compared to the previous salinity level.

**Figure. 1.**
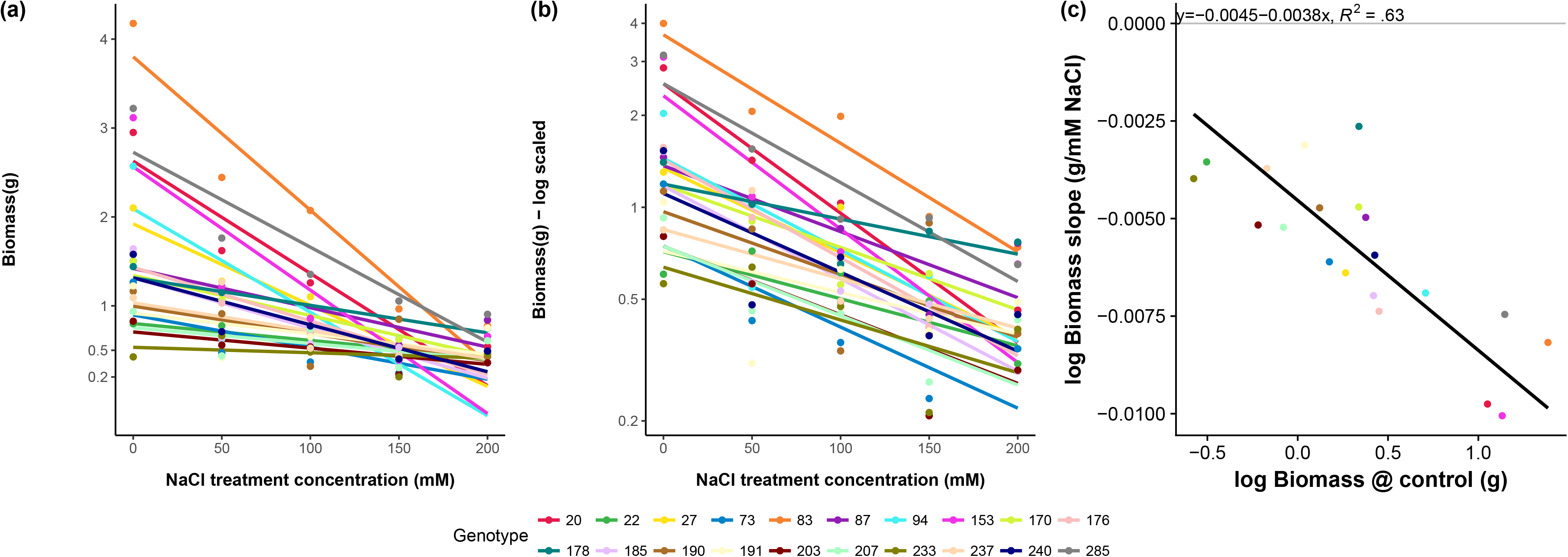
Biomass response of twenty sunflower genotypes to five salinity treatments. Points indicate genotype estimated marginal means of **(a)** absolute biomass, showing a steeper decline for more vigorous gentoypes under control conditions. After **(b)** natural log transformed total plant biomass in a salinity treatment, proportional declines are much more similar. Lines are fitted slopes per genotype from the mixed effects model incorporating the split plot design. Acros vigour at control conditions **(c)** standardized major axis regression shows a significant, negative, relationship between biomass at zero mM NaCl and the proportional effect of increased salinity (slope of log transformed biomass to salinity).

However, when explicitly taking vigour (biomass under the control treatment) into account we found a strong (p<0.001, R^2^>0.63) correlation between vigour and the proportional effect of increased salinity (slope in figure 1b) on biomass **(Figure 1c)**. Genotypes with greater biomass at zero mM NaCl had a greater proportional decrease in biomass under saline conditions. Other plant vigour indicators, height and stem diameter, showed comparable results **(Supplemental Figure 2 and 3)**. Overall, high vigour was correlated with a stronger effect (i.e. decrease in growth) of increased salinity.

Seedling survival decreased with increasing salinity (p<0.05). Survival rates were high under 0, 50 and 100 mM NaCl but rapidly declined to median 50% surviving individuals per genotype at 200 mM NaCl. While only 3 out of 20 genotypes had complete mortality under 200 mM NaCl, we did not detect significant differences in mortality among genotypes, likely owing to relatively small sample size for assessing mortality within genotypes **(Supplemental Fig. 4)**. However, when pooled across genotypes, these results demonstrate these cultivated sunflower genotypes were moderately salt tolerant with only limited death up to 100/150 mM NaCl.

Leaf elemental concentrations were affected by salinity level. Of the twelve elements measured, six either had a significant effect of treatment or a significant interaction between genotype and treatment **(Table 1)**. With increasing salinity, there were strong increases in Na concentration and K:Na ratio, coupled with decreases in both S and K **(Figure 3)**. Genotypes that had a shallow slope of leaf Na increase with increasing salt treatment also had a shallow slope of leaf K decrease. The response of other elements (N, P, Ca, Mg, Mn, Fe, Cu, Zn, B) to increased salinity can be seen in **Supplemental figure 5**.

**Figure. 2.**
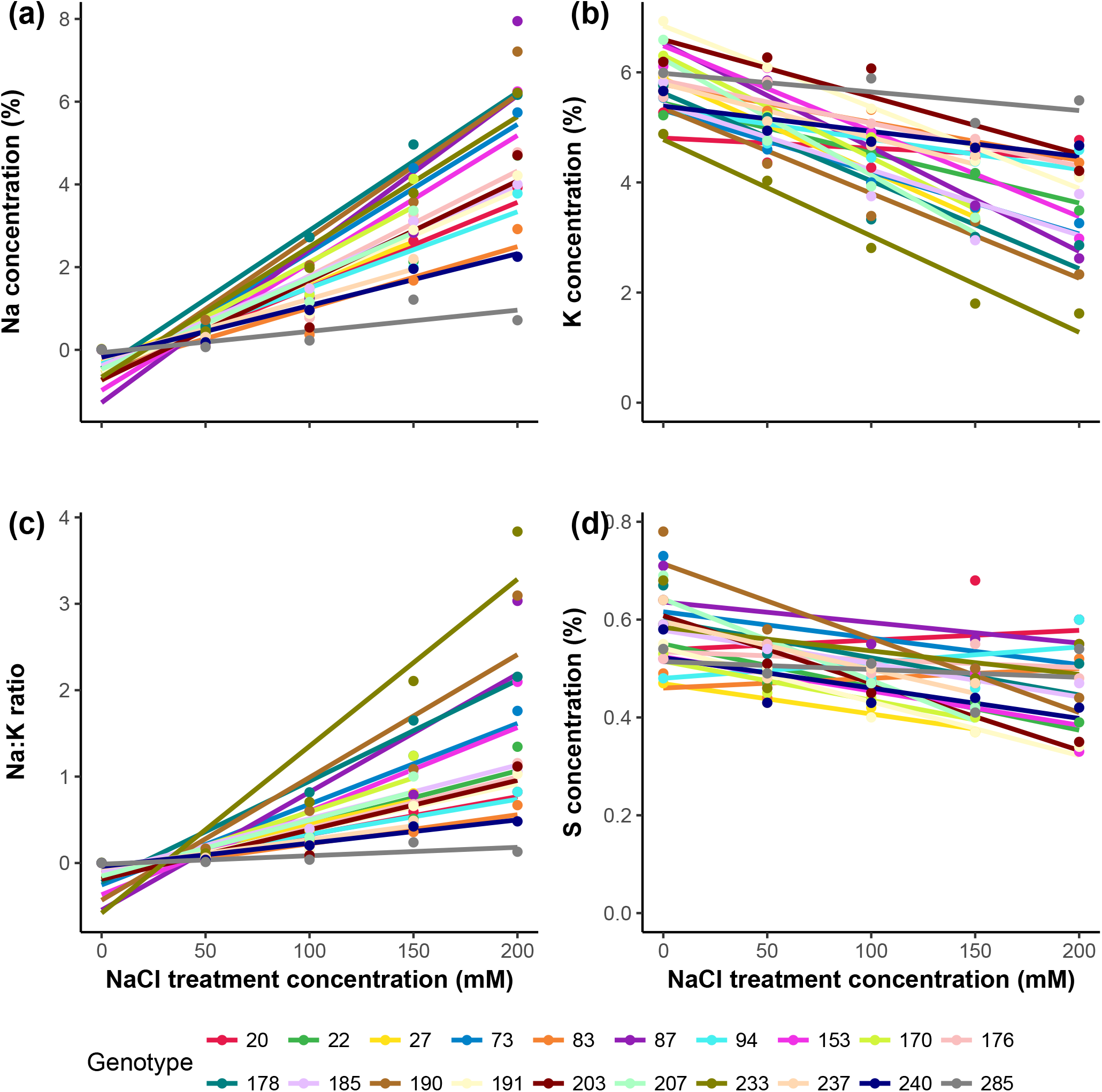
Shifts in leaf tissue major element concentration with increasing salinity treatment. Points indicate tissue element concentration (mass %) of bulked (n=1-4) ground and homogenized leaf tissue at five soil salinity concentrations for twenty sunflower genotypes. Lines are fitted linear regression per genotype across all five salinity levels. **(a)** leaf sodium, Na, concentration, **(b)** leaf potassium, K, concentration, **(c)** Ratio of leaf sodium to leaf potassium, K:Na, **(d)** leaf sulfur, S, concentration.

**Figure. 3.**
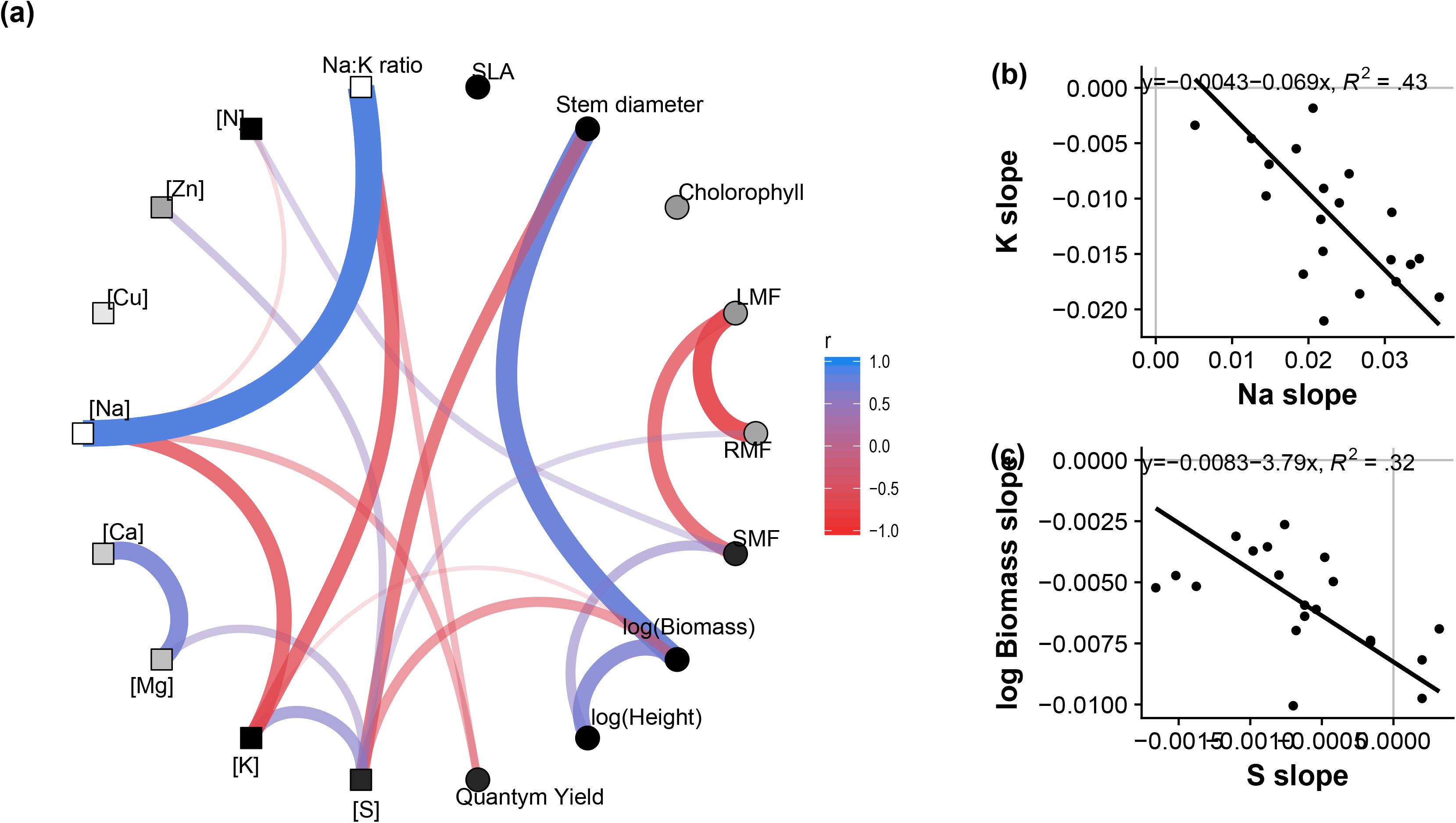
Correlated trait shifts in response to increasing salinity treatment. **(a)** Correlation across twenty sunflower genotypes of the slopes in genotype trait adjustment to salinity increases. Genotype slopes were calculated from mixed model for harvest traits (circles) and linear regression for elemental composition (squares). Correlation among trait slopes were calculated using standardized major axis (SMA) regresion. Edges are colored and scaled by correlation sign and strength. Nodes are colored by sign of slope to salinity, black: all genotypes negative, white: all genotypes positive, grey: mixed slopes across genotypes. Only edges with r>0.4 and p<0.05 are shown. Examples of trait correlations. **(b)** Genotypes slope of Na increase with increasing salinity vs genotypes slope of K decrease with increasing salinity. **(c)** Genotypes slope of sulfur concentration decrease vs genotypes slope of log biomass decrease.

### Correlated trait adjustments to increasing salinity

In the higher salinity treatments, leaf Na concentration was negatively correlated with leaf K and sometimes leaf N. Additionally, leaf Ca, Mg and Zn levels were positively correlated with each other. In all salinity treatments, plant morphological traits (height, allocation, stem diameter) and biomass traits were highly correlated (**Supplemental figure 6**). However, the correlation between SLA and chlorophyll content was only evident in lower salinity treatments. Significant correlations among elemental composition and morphological and physiological traits did not follow consistent patterns across all salinity treatments, as different traits were correlated under different salinity treatments.

We also examined correlations between traits’ slopes to increasing salinity (**Figure 4a**). For a complete graph of slopes correlation, see **Supplemental figure 7**. In terms of elemental composition, there was a relationship between leaf Na, K and N such that genotypes that had a steeper slope of leaf Na increase with increasing salinity having had a steeper slope of K (**Figure 4b**) and N decrease with increasing salinity. Leaf K:Na ratio was negatively correlated with quantum yield, both dark adapted and in the light. Additionally, the slope in leaf S concentration decrease was correlated with the slope of leaf P, K, Mg, and Zn concentration decrease. While there were differences among genotypes in leaf Na accumulation and K:Na ratio, these factors were not linked to differences in the effect of salinity on biomass. Rather, only the effects of salinity on genotype leaf S (**Figure 4c**) and K concentrations were correlated with the effect of salinity on biomass. Surprisingly, a greater reduction in leaf S and K concentration was correlated with a lower effect of salinity on biomass.

## Discussion

Salinity is a key stress limiting agricultural productivity worldwide. In this study we grew twenty inbred sunflower genotypes under five salinity levels (0, 50, 100,150, 200 mM NaCl) for three weeks and determined the effect of salinity on biomass, leaf elemental composition and leaf trait correlations. Despite strong survival, we found that increasing soil salinity had a strong, proportional to biomass, effect on biomass accumulation. For every 50 mM increase in soil NaCl, genotypes exhibited an average 25% decrease in total biomass compared to the previous lower salinity level. This greater proportional decline was associated with decreased leaf potassium and sulfur concentrations. However, within this average decrease, we found that the more vigorous genotypes under control conditions had a greater proportional decrease in biomass. More vigorous genotypes at control were less tolerant to salinity. Multiple mechanisms could lie at the heart of this difference in tolerance. Before looking at costs underlying this vigour/tolerance trade-off first we need to find evidence for what kind of tolerance mechanisms these sunflower genotypes employ.

One crucial factor in salt tolerance is the maintenance of leaf (specifically high cytosol) K:Na ratio (Shabala & Cuin, 2008; Shahbaz et al., 2011). Across salinity levels, we found whole leaf K:Na ratio to be strongly impacted by accumulation of Na^+^ and reduced concentrations of K^+^ (**Figure 4a**). Surprisingly, the maintenance of whole leaf K:Na ratio was not correlated with the effect of salinity on biomass. Suggesting that maintenance of leaf K:Na ratio is less important for sunflower salinity tolerance. However, a mechanism underlying this discrepancy could be that tolerant genotypes sequester excess sodium in the vacuole, thereby maintaining cytosolic K:Na ratio while whole leaf K:Na ratio altered (Bassil, Zhang, Gong, Tajima, & Blumwald, 2019; Blumwald, 2000). The lack of a connection of tolerance and whole leaf K:Na ratio suggests this vacuolar sequestration mechanisms to be important, as in other species (Hasegawa, 2013).

Contrary to the generally expected results, a greater reduction in leaf S and K concentration was correlated to a lower effect of salinity on biomass (Nazar, Iqbal, Masood, Syeed, & Khan, 2011; Shabala & Cuin, 2008). While running counter to the idea that K^+^ retention is key in salinity tolerance, these results do fit within a relatively novel understanding of the role of K^+^ in salinity tolerance. K^+^ loss has been hypothesized as a metabolic switch transitioning cell energy budgets from metabolic processes to stress defense and repair (Demidchik, 2014). Under salt stress, rebalancing a cell’s diminished energy pool could be key in preventing cell death (Shabala, 2017). Alternatively, decreasing leaf level K^+^ and S concentrations could be indicative of a redistribution of these elements to the roots. Indeed, across seven crop species, maintenance of root K^+^ concentrations has been shown to be negatively correlated with shoot K^+^ concentration (Wu, Zhang, Giraldo, & Shabala, 2018), and root sulfur content in the form of sulpholipids has been linked to salt tolerance (Erdei, Stuiver, & Kuiper, 1980; Stuiver, Kuiper, Marschner, & Kylin, 1981). These results suggest K^+^ and S to play a role in salinity tolerance, but likely not in the leaf.

Costs associated with these sequestration, K^+^ and S redistribution, and other mechanisms related to the ionic and osmotic effects of salinity stress could lie at the heart of the apparent trade-off between vigour under control conditions and tolerance to salinity. For example, strong root membrane potentials via constitutive high levels of H^+^ pumping serves to exclude Na^+^ out from the root system (Bose et al., 2015; Chen et al., 2007) transporters can pump sodium out of the xylem back into the roots/stem (Munns & Tester, 2008; Schroeder et al., 2013), and/or into the vacuole to sequester it safely (Hasegawa, 2013). However, these molecular mechanisms come at a metabolic cost that may result in decreased vigour under benign conditions. Additionally, genotypes with a low stomatal conductance could conserve water better under osmotically stressed conditions (Munns & Tester, 2008). This does come at the cost of reduced carbon uptake and nutrient flow from soil to plant, diminishing vigour. As we found no link between tolerance and whole leaf sodium concentration these results suggest that the cost of sodium exclusion at the roots alone isn’t enough to explain the vigour/tolerance trade-off. Rather costs potentially associated with reduced stomatal conductance and/or sequestering of sodium in the vacuole and/or other organs are likely both important as a factor underlying the trade-off between plant vigour and /salt tolerance trade-off.

In contrast to safflower (Yeilaghi et al., 2015) and other longer duration sunflower experiments (Gabriel Ceccoli et al., 2012; Di Caterina, Giuliani, Rotunno, De Caro, & Flagella, 2007; Shahbaz et al., 2011), we found no link between leaf Na and tolerance. A key difference in these other studies and our experiment is the duration of stress. Potentially, when subjected to longer periods of stress, the ionic component, toxic Na^+^, of salt stress (Munns & Tester, 2008) will become stronger. Combined with our results, this suggest a switch from tolerance to the osmotic component of salt stress being dominant in early vegetative stages to tolerance to the ionic component of salt stress being dominant during reproductive stages. Taking both of these components and ontogenetic shifts into account will be important for improving sunflower salt tolerance at all life stages.

While our results provide some indication for mechanisms used for salt tolerance in cultivated sunflower more detailed work is needed to disentangle these. As we found whole leaf elemental composition to be only moderately linked to salt tolerance exploring Na sequestration in the vacuole as well as a whole plant elemental budget approach is needed to shed light on the role of leaf, stem, and root elemental content and salt tolerance. Furthermore, exploring strategies to ameliorate the effect of salinity, such as exogenous application of compatible solutes and cations (Shabala & Cuin, 2008), the magnitude of osmotic adjustment (Serraj & Sinclair, 2002), and the extent of sodium sequestration versus sodium exclusion (Munns & Tester, 2008), will provide exciting areas for further research in sunflower.

Here we defined tolerance as a low proportional effect of stress on biomass. However, several competing definitions of tolerance could lead to different interpretations of the results. For instance, while there is a clear trade-off between plant vigour (biomass at zero mM NaCl) and the proportional decrease in biomass due to salinity, the genotypes that were most vigorous still maintained highest biomass under stressed conditions **(Supplemental figure 8)**. Thus, an argument could be made that the genotypes that are most vigorous are inherently the most tolerant, and they will always perform well, even under stressful growing conditions. However, from an agricultural standpoint, the ideal genotype would have both high vigour as well as a low proportional decrease in biomass under stress. This would ensure that crops would maintain high yields (i.e. vigour, growth) under stressful growing conditions. Given that, ideal candidates for future work would be genotypes that are more tolerant than expected, or above the fitted “expected” line in figure 1c (genotype 178, for example). Studying **specifically** these genotypes will allow us not only to identify the traits associated with stress tolerance, but also to more thoroughly investigate the traits and mechanisms that allow for greater than expected stress tolerance.

The strong survival across twenty genotypes up to 100 mM NaCl demonstrates moderate salt tolerance for these genotypes, consistent with the crop as a whole (**Supplemental figure 1**; (Katerji et al., 2000)). As it is not only the growth of surviving plants that matters for crop yield but also the establishment and survival of seedlings (Flowers, 2004), this suggests that with limited overplanting, sunflower could be a suitable crop for salinized soils. Given their capacity to hybridize with closely related, more salt tolerant sunflower species (Rosenthal, Schwarzbach, Donovan, Raymond, & Rieseberg, 2002) there is a high potential for incorporation of beneficial traits to boost their survival and growth under saline conditions.

A concern with growing crops under stressful conditions is whether the highest yielding genotypes under benign conditions are most suitable for growth under stressful conditions, since high yield could come at the penalty of reduced stress tolerance. Results here suggest that for sunflower under saline conditions, there is a trade-off between vigour and a higher proportional effect of increased salinity on biomass. However, this increased effect of salinity is not big enough to result in a complete reversal of genotype performance, because genotypes with higher vigour also maintained larger biomass under saline conditions. In order to increase the yield of genotypes under saline conditions, traits and genes that confer stress tolerance in the tolerant genotypes need to be bred into high yielding varieties. Genotypes that exhibit different K:Na ratios, and the intriguing correlation between sulfur concentration and the proportional effect of salinity on biomass, suggests there is ample variation in leaf traits that could be explored to further improve sunflower salt tolerance. Testing the effect of increased salinity on a larger diversity panel of sunflower genotypes (Mandel et al., 2011; Nambeesan et al., 2015) will reveal the extent of variation in these traits as well as target genomic regions linked to salinity tolerance.

## Acknowledgements

This work was financially supported by grant NSF1444522 to LAD. We would like to thank K. Bettinger, K. Davis, M. Boyd, K. Tanner, A. Kerr, T. Nortier, L. Ceelen and Y. Ceelen for their help during measurements and harvest. We would like to thank the reviewers for their time and insightful comments on the manuscript.

