## Supplemental Table 1 for "Vigour/tolerance trade-off in cultivated sunflower (*Helianthus annuus*) response to salinity stress is linked to leaf elemental composition"

**Supplemental Table 1.** List of genotypes used in this study, it’s common name, the corresponding plant ID from the USDA GRIN Database for each genotype, and the market type of each genotype.

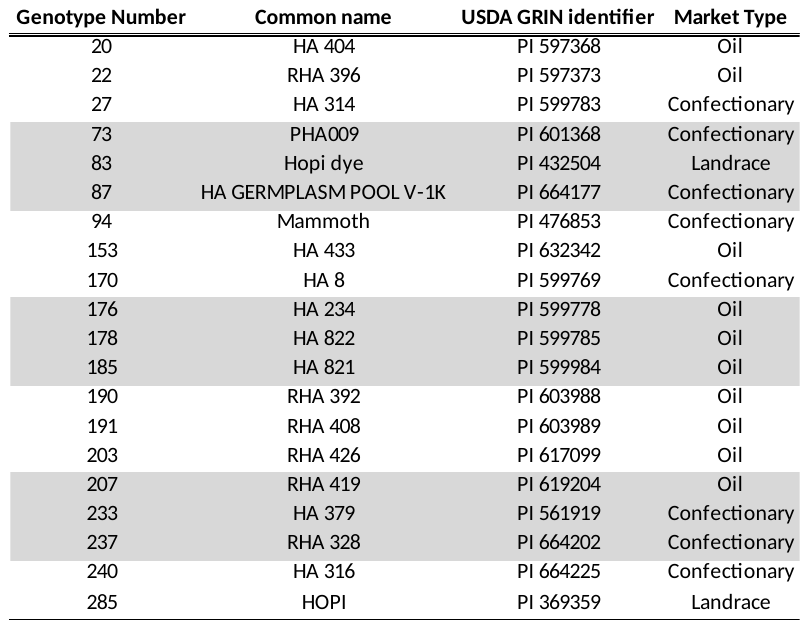

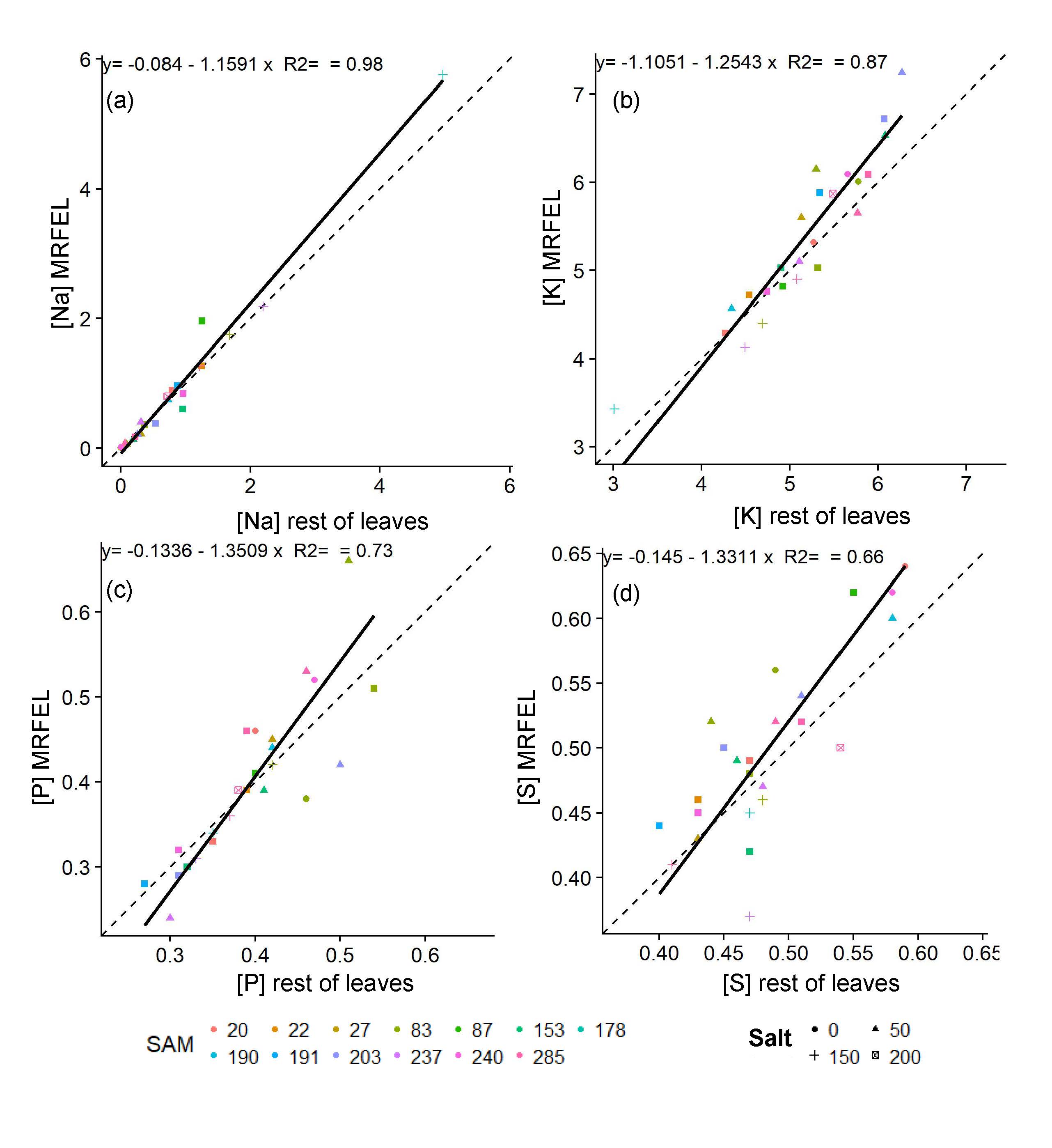

**Supplemental Figure 1. Most recent fully expanded leaf element concentration versus rest of leaves element concentration.** Correlations between element concentration in the most recent fully expanded leaf (MRFEL) and the rest of the leaf tissue for (a) sodium, Na, (b) potassium, K, (c) phosphorous, P, and (d) sulfur, S.

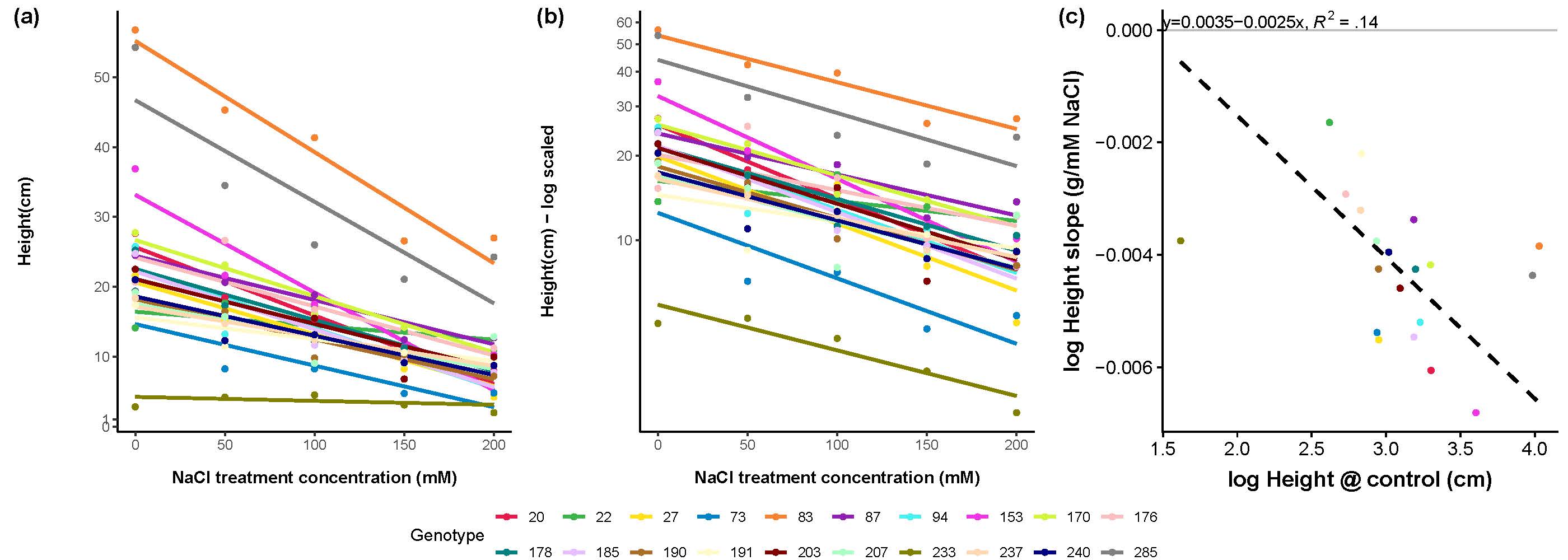

**Supplemental Figure 2. Height response of twenty sunflower genotypes to five salinity treatments.** Points indicate genotype estimated marginal means of **(a)** absolute height and **(b)** natural log transformed height in a salinity treatment. Lines are fitted slopes per genotype from the mixed effects model incorporating the split plot design. (**c**) Standardized major axis regression of height at zero mM NaCl and the proportional effect of increased salinity (slope of log transformed height to salinity).

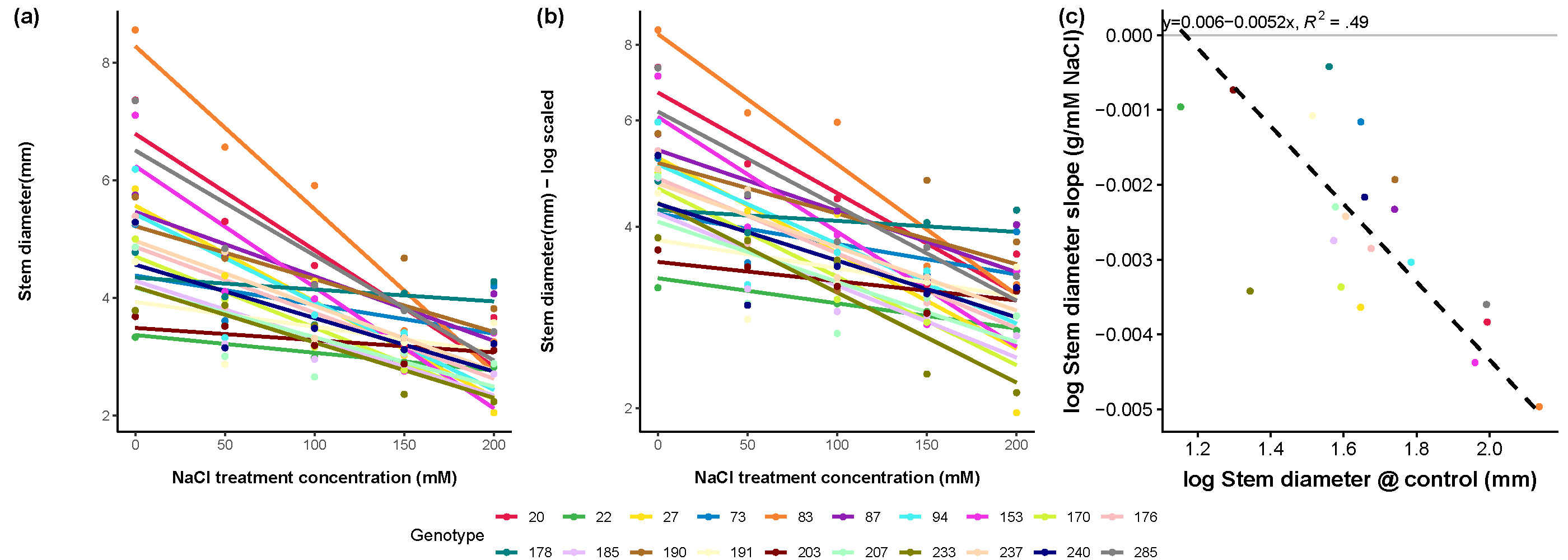

**Supplemental Figure 3. Stem Diameter response of twenty sunflower genotypes to five salinity treatments.** Points indicate genotype estimated marginal means of **(a)** absolute stem diameter and **(b)** natural log transformed stem diameter in a salinity treatment. Lines are fitted slopes per genotype from the mixed effects model incorporating the split plot design. (**c**) Standardized major axis regression of stem diameter at zero mM NaCl and the proportional effect of increased salinity (slope of log transformed stem diameter to salinity).

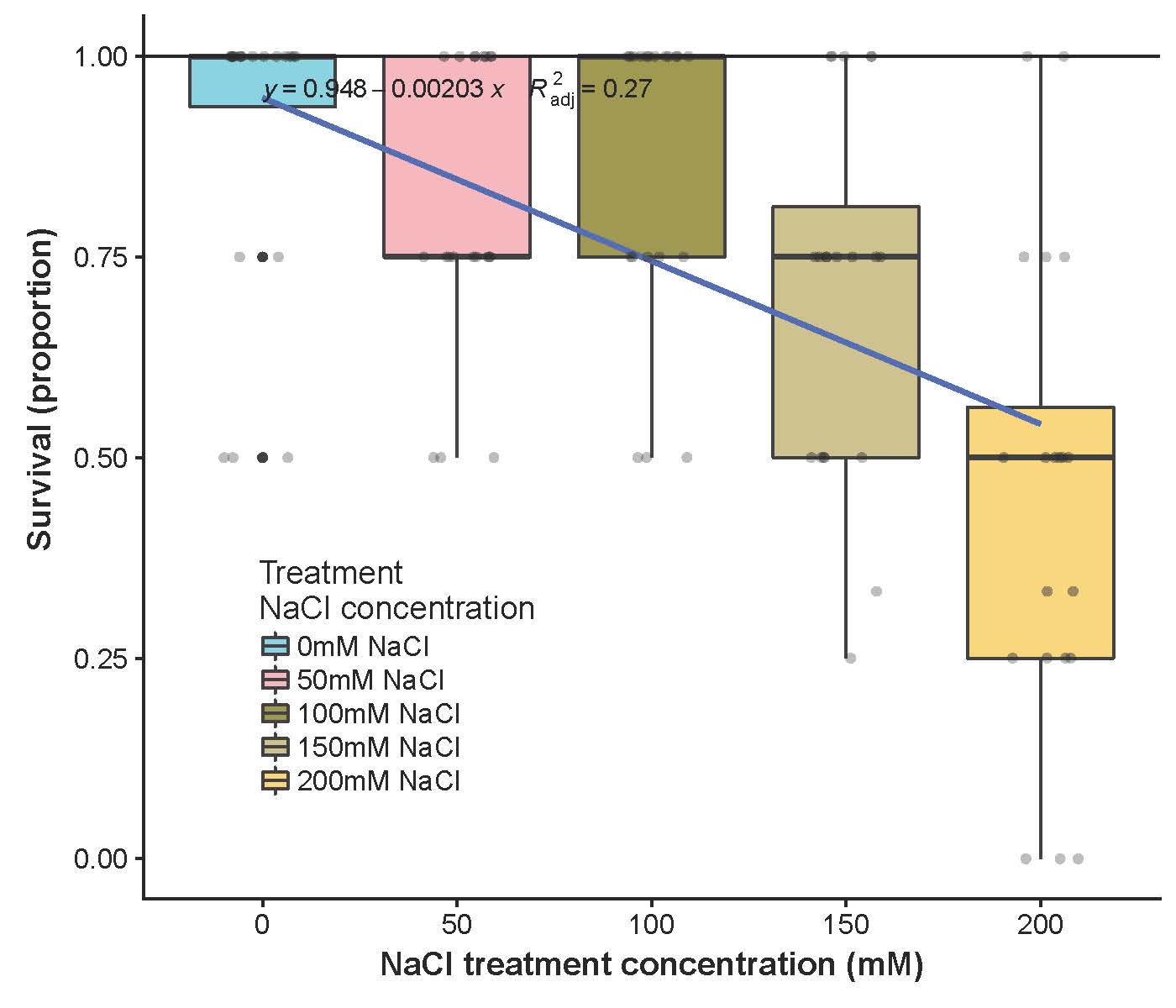

**Supplemental Figure 4. Sunflower survival under salinity stress.** Points indicate proportional survival of a given genotype (n=3-4 per genotype) at a given salinity concentration. Boxplots indicate distribution of survival of twenty genotypes. The fitted line indicates aggregate survival across salinity concentrations.

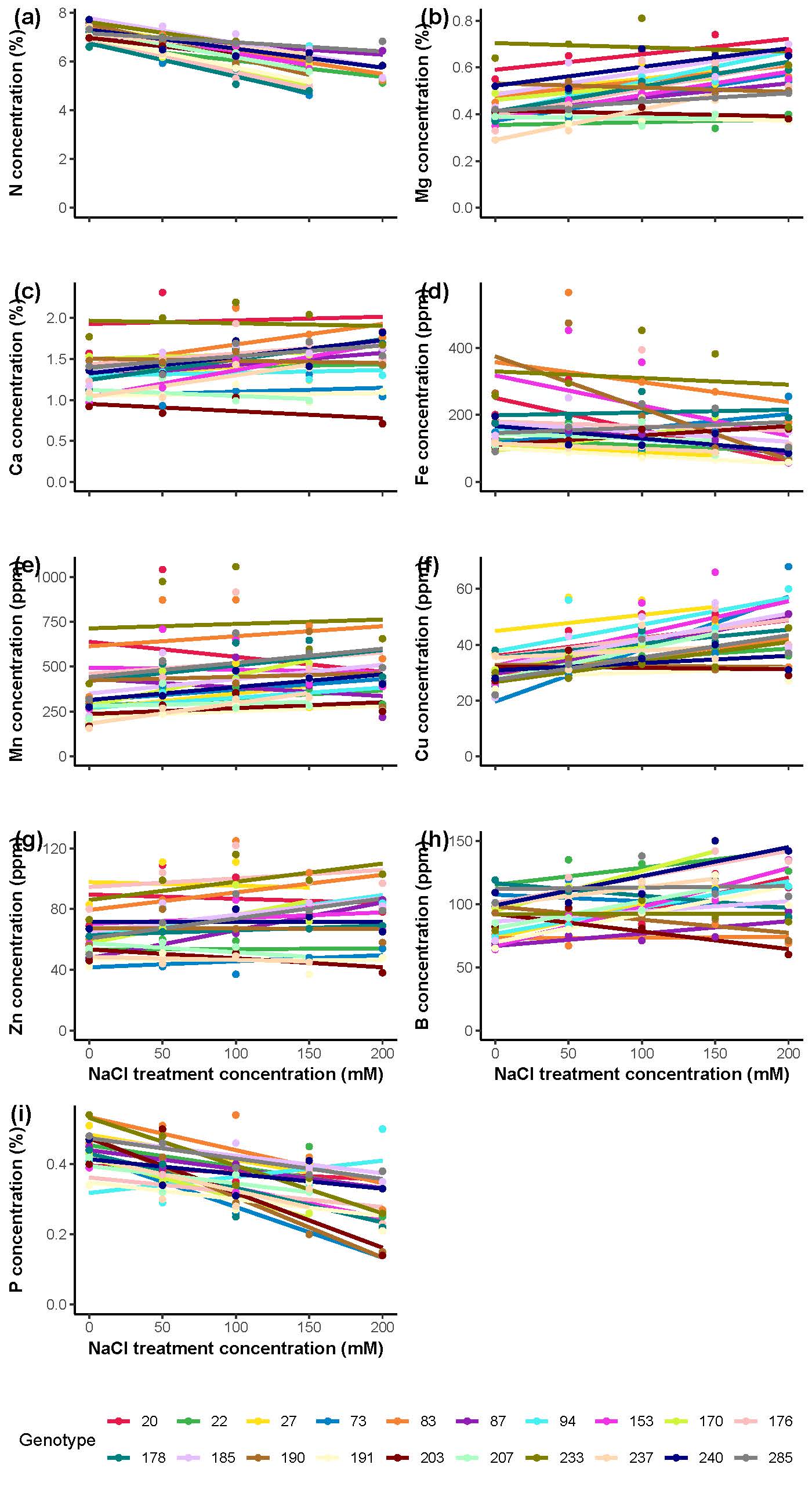

**Supplemental Figure 5. Shifts in leaf tissue element composition under salinity stress.** Points indicate leaf tissue element concentration (mass %) of bulked (n=1-4) ground and homogenized leaf tissue at five soil salinity concentrations for twenty sunflower genotypes. Lines are fitted linear regression per genotype across all five salinity concentrations. **(a)** sulfur, S, concentration, **(b)** magnesium, Mg, concentration, **(c)** calcium, Ca, concentration, **(d)** iron, Fe, concentration, **(e)** manganese, Mn, concentration, **(f)** copper, Cu, concentration, **(g)** zinc, Zn, concentration, **(h)** boron, B, concentration **(i)** phosphorus, P, concentration.

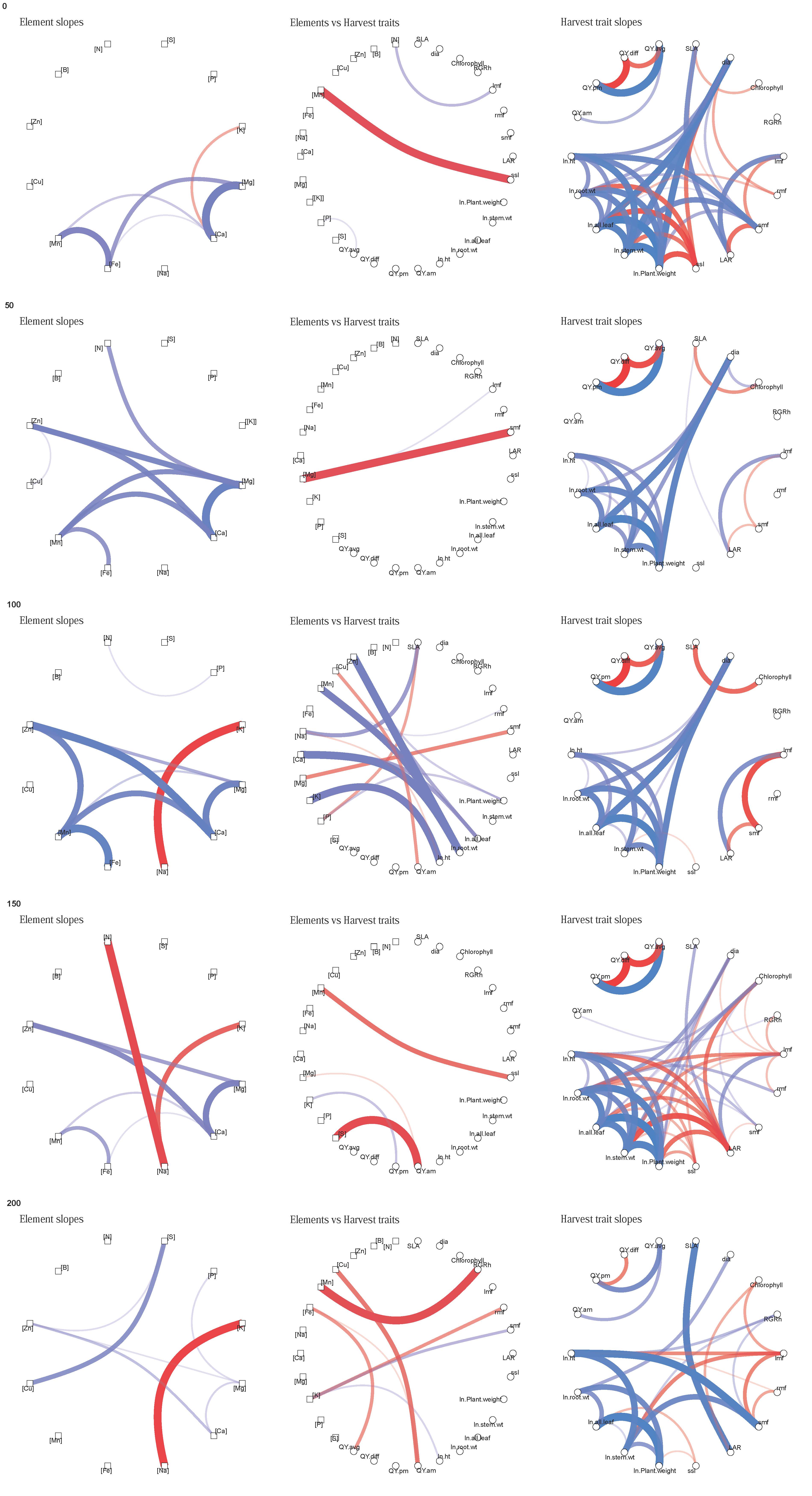

**Supplemental Figure 6. Trait correlation networks at five salinity concentrations.** Correlation across twenty sunflower genotypes of trait values at five salinity concentrations (0mM, 50mM, 100mM, 150mM, and 200mM NaCl). **(a)** Correlation network of element composition. **(c)** Correlation network of traits measured at harvest. **(b)** Correlation network only showing the connections between elements and harvest traits. Edges are colored and scaled by correlation strength. Only edges with r>0.4 and p<0.05 are shown.

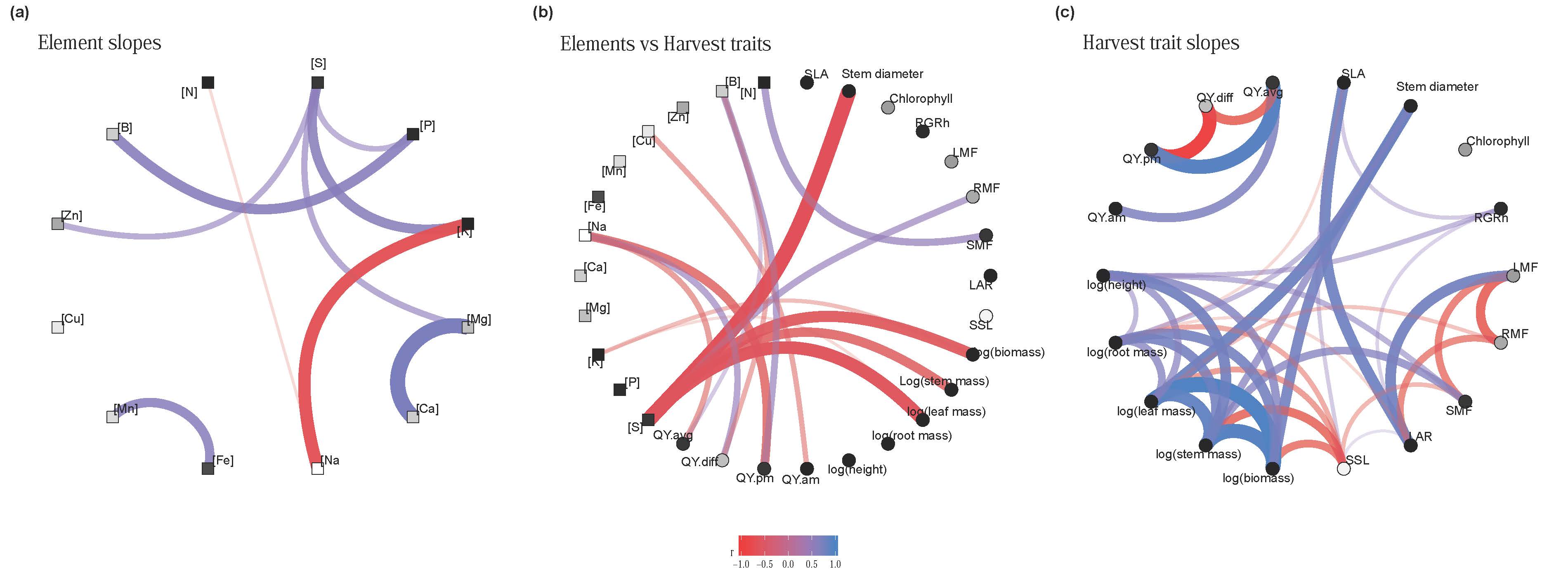

**Supplemental Figure 7. Correlated trait shifts across salinity concentrations.** Correlation across twenty sunflower genotypes of the slopes in genotype trait adjustment to salinity increases. Genotype slopes were calculated from mixed model for harvest traits and linear regression for elemental composition. **(a)** Correlation network of responses to salinity in element composition. **(c)** Correlation network of responses to salinity in traits measured at harvest. **(b)** Correlation network only showing the connections between elements and harvest traits. Edges are colored and scaled by correlation sign and strength. Nodes are colored by sign of slope to salinity, black: all genotypes negative, white: all genotypes positive, grey: mixed slopes across genotypes. Only edges with r>0.4 and p<0.05 are shown.

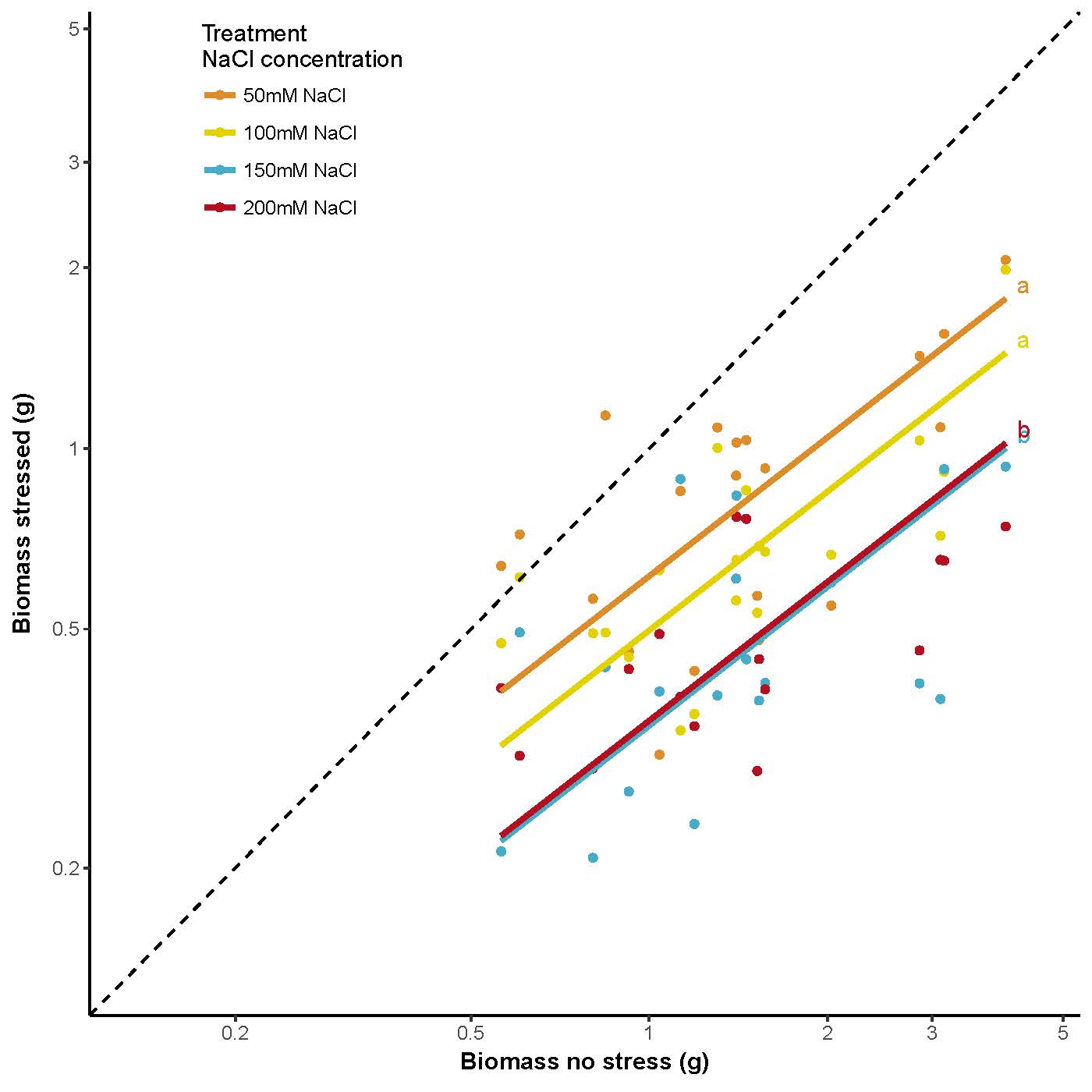

**Supplemental Figure 8. Relationship between biomass at no stress, and salinity stressed conditions.** The dotted line represents the 1:1 line indicating no effect of salinity stress. Note that both axes are natural log transformed, indicating that a similar distance from the 1:1 line is a similar *proportional* decrease in biomass. Points indicate genotype estimated marginal means of natural log transformed total plant weight at a given soil salinity concentration. Lines are fitted lines from standardized major axis regression across genotypes per salinity stress treatment. Different letters indicate significantly different intercepts.
